# Prospecting for zoonotic pathogens using targeted DNA enrichment

**DOI:** 10.1101/2022.12.06.519193

**Authors:** Egie E. Enabulele, Winka Le Clec’h, Emma K. Roberts, Cody W. Thompson, Molly M. McDonough, Adam W. Ferguson, Robert D. Bradley, Timothy J. C. Anderson, Roy N. Platt

## Abstract

There are over 60 zoonoses linked to small mammals, including some of the most devastating pathogens in human history. Meanwhile, millions of museum-archived tissues are available to understand natural history of these pathogens. Our goal is to maximize the value of museum collections for pathogen-based research using targeted sequence capture. To this end, we have generated a probe panel that includes 39,916, 80bp RNA probes targeting 32 pathogen groups, including bacteria, helminths, fungi, and protozoans. Lab generated, mock control samples show that we are capable of enriching targeted loci from pathogen DNA 2,882 to 6,746-fold. Further, we were able to identify bacterial species in museum-archived samples, including *Bartonella*, a known human zoonosis. These results show that probe-based enrichment of pathogens is a highly customizable and efficient method for identifying pathogens from museum-archived tissues.

## Introduction

Many important human pathogens result from zoonotic transmission. These include 61% of known human pathogens, and 75% of emerging human pathogens (1). For example, Rabies virus is transmitted via the saliva of infected animals (2). The plague bacteria (*Yersina pestis*), the causative agent of the largest documented pandemic in human history, which reduced the population of Europe by 30-50%, was transmitted from rats to humans via fleas (3). Other zoonoses include Ebola virus (4), tularemia (*Francisella tularensis*) (5), and tuberculosis (6). The SARS-CoV-2 pandemic, thought to have a bat reservoir, has stimulated renewed emphasis on zoonotic pathogen surveillance (7, 8).

Natural history museums are repositories of biological information in the form of archived voucher specimens that represent a significant and underutilized resource for studying zoonotic pathogens (9-13). Originally, specimens were archived as dried, skin and skeletal vouchers or preserved in fluids (ethanol) after fixation with formalin or formaldehyde. Now, best practices include preserving specimens and associated soft-tissues in liquid nitrogen (−190°C) or mechanical freezers (−80°C) from the time they are collected (14). These advances in preservation make it possible to extract high-quality DNA and RNA that can be used for pathogen surveillance. For example, retroactive sampling of museum archived tissues from the American Southwest found that Sin Nombre Virus, a new world hantavirus, was circulating in wild rodent populations almost 20 years before the first human cases were reported (15).

It is critical to develop a range of tools for extracting pathogen information from museum archived samples. Targeted sequencing via probe enrichment has become the tool of choice for human medical genomics (16), population genetics (ex. 17), phylogenetics (18), and ancient DNA. These methods are designed to enrich very small amounts of DNA target from a background of contaminating DNA. Further, Vernot *et al*. (19) were able to sequence Neanderthal DNA by targeted enrichment from cave sediment. Probe-based, targeted sequencing has been used to enrich pathogens from complex host:pathogen DNA mixtures (20). For example, Keller *et al*. used probes to capture and sequence complete *Yersinia pestis* genomes from burial sites more 1,500 years old (21). Enrichment is frequently achieved by designing a panel of probes to specifically target a handful of pathogens of interest (22, 23). Similarly, commercial probe sets are available for many types of viruses and human pathogens (22-24). However, many of these probe sets are limited to specific pathogens which may not impact other host species.

Our goal was to develop a panel of biotinylated baits, or probes, to identify the eukaryotic and bacterial pathogens responsible for 32 major zoonoses (Table 1). We aimed to capture both known and related pathogens, utilizing the fact that probes can capture sequences that are ≤10% divergent. To do this we used a modified version of the ultra-conserved element (UCE) targeted sequencing technique (25, 26) to specifically enrich pathogen DNA. Briefly, biotinylated baits are designed to target conserved genomic regions among diverse groups of pathogens. These baits are hybridized to a library potentially containing pathogen DNA. Bait-bound DNA fragments are enriched during a magnetic bead purification step prior to sequencing. The final library contains hundreds or thousands of orthologous loci with single nucleotide variants (SNVs) or indels from the targeted pathogen groups that can then be used for population or phylogenetic analyses.

**Table 1.** Target Zoonoses.

| Pathogen group | Taxonomic level | Focal Pathogen | Ex. Zoonoses |
| --- | --- | --- | --- |
| Anaplasma | Genus | *Anaplasma phagocytophilum* | Anaplasmosis |
| Apicomplexa | Phylum | *Plasmodium falciparum* | Malaria |
| Bacillus cereus group* | Species Group | *Bacillus anthracis* | Anthrax |
| Bartonella | Genus | *Bartonella bacilliformis* | Bartonellosis; Cat-Scratch Fever |
| Borrelia | Genus | *Borrelia burgdorferi* | Lyme Disease; Relapsing Fever |
| Burkholderia | Genus | *Burkholderia mallei* | Glanders |
| Campylobacter | Genus | *Campylobacter jejuni* | Campylobacteriosis |
| Cestoda | Class | *Taenia multiceps* | Taeniasis |
| Chlamydia | Genus | *Chlamydia trachomatis* | Chlamydia |
| Coxiella | Genus | *Coxiella burnetii* | Q Fever |
| Ehrlichia | Genus | *Ehrlichia canis* | Ehrlichiosis |
| Eurotiales | Order | *Talaromyces marneffeii* | Talaromycosis |
| Francisella | Genus | *Francisella tularensis* | Tularemia |
| Hexamitidae | Family | *Giardia intestinalis* | Giardiasis |
| Kinetoplastea | Class | *Leishmania major* | Leishmaniasis |
| Leptospira | Genus | *Leptospira interrogans* | Leptospirosis |
| Listeria | Genus | *Listeria monocytogenes* | Listeriaosis |
| Mycobacterium | Genus | *Mycobacterium tuberculosis* | Tuberculosis; Mycobacteriosis |
| Nematodes (clade I) | Phylum (Clade) | *Trichinella spiralis* | Trichinosis |
| Nematodes (clade III) | Phylum (Clade) | *Brugia malayi* | Filariasis |
| Nematodes (clade IVa) | Phylum (Clade) | *Strongyloides stercoralis* | Strongyloidiasis |
| Nematodes (clade IVb) | Phylum (Clade) | *Steinernema carpocapsae* |  |
| Nematodes (clade V) | Phylum (Clade) | *Haemonchus contortus* |  |
| Onygenales | Order | *Histoplasma capsulatum* | Histoplasmosis |
| Pasteurella | Genus | *Pasteurella multocida* | Pasteurellosis |
| Rickettsia | Genus | *Rickettsia rickettsii* | Typhus; Spotted Fevers |
| Salmonella | Genus | *Salmonella enterica* | Salmonellosis |
| Streptobacillus | Genus | *Streptobacillus moniliformis* | Rat-Bite Fever |
| Trematoda | Class | *Schistosoma mansoni* | Schistosomiasis; Fascioliasis |
| Tremellales | Order | *Cryptococcus neoformans* | Cryptococcosis |
| Trypanosoma* | Genus | *Trypanosoma cruzi* | Sleeping Sickness; Chagas disease |
| Yersinia | Genus | *Yersinia pestis* | Plague |

\*supplemented with additional probes/baits

## Methods

A detailed description of the methods is provided in a supplemental file and at protocols.io (DOI: dx.doi.org/10.17504/protocols.io.5jyl8jnzrg2w/v1). A summary of the methods is provided below and in Figures 1-3. Code is available at https://www.github.com/nealplatt/pathogen_probes (DOI: 10.5281/zenodo.7319915). Raw sequence data is available under the NCBI BioProject ID PRJNA901509.

**Figure 1.**
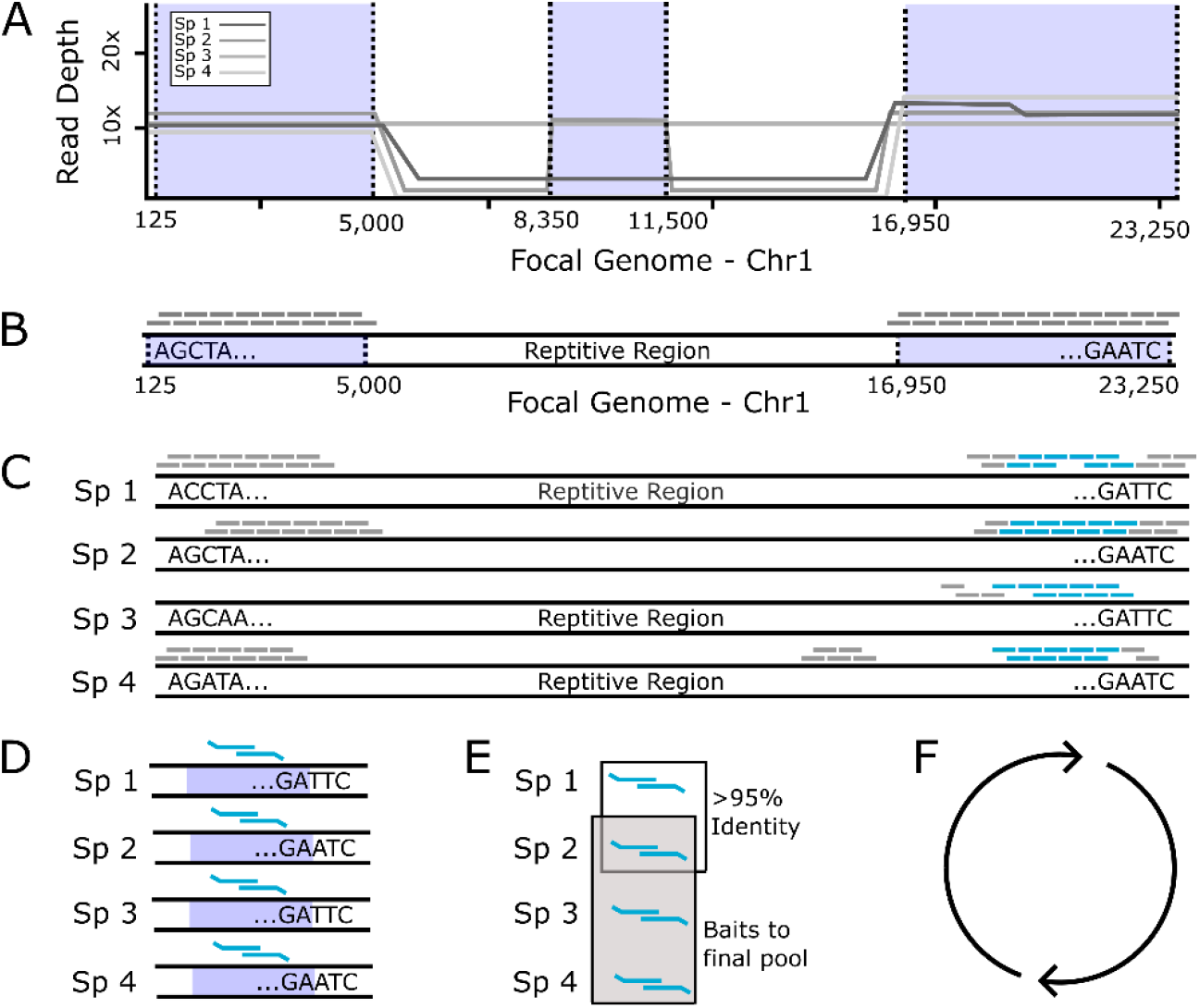
Designing a probe panel. (A) Simulated reads from each pathogen within a group were mapped back to a single focal genome. (B) Next, we identified regions with consistent coverage from each member of the pathogen group to identify putatitve, orthologous loci and generated a set of in-silico probes from the focal genome. (C) These in-silico probes were then mapped back to the genomes of each member in the pathogen group to find single copy, orthologous regions, present in a majority of members. (D) Next, we designed two, overlapping 80bp baits to target these loci in each member of the pathogen group and (E) compared them to one another to remove highly similar probes. One probe was retained from each group of probes with high sequence similarity (>95%). Then, we identified the probes necessary to capture 49 loci in that pathogen group. (F) This process was repeated for the next pathogen group. Finally, all probes were combined together into a single panel.

**Figure 2.**
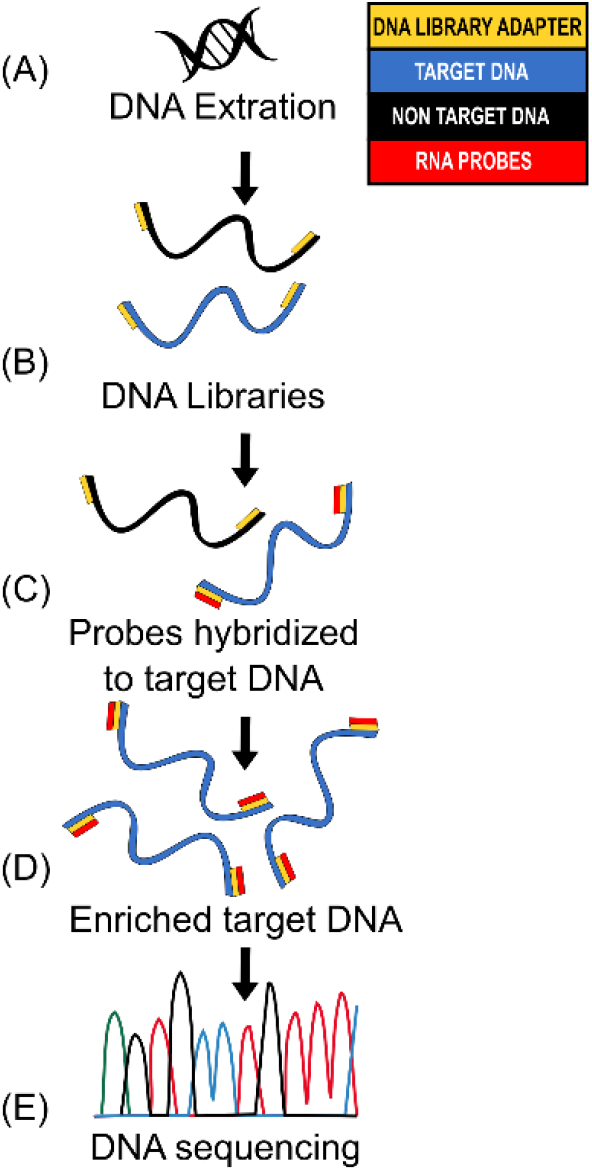
Targeted DNA capture workflow overview. (A) Genomic DNA extrated using Qiagen Dneasy kit (B) NGS libraries prepared using KAPA Hyperplus kit and barcoding each library with IDT xGen Stubby Adapter-UDI Primers (C) RNA probes hybridization using the high sensitivity protocol of myBaits® v.5 (D) 15 cycles PCR amplification of enriched libraries (E) Libraries sequenced on an Illumina Hi-Seq 2500 platform.

### Panel development

We developed a panel of biotinylated baits for the targeted sequencing of 32 zoonotic pathogens. To do this, we used Phyluce v.1.7.1 (25, 26) protocol to design baits for conserved loci within each pathogen group. First, we simulated and mapped reads from each species within a pathogen group to a focal genome assembly (Figure 1A; Supplemental Table 1). We used the mapped reads to identify putative orthologous loci that were >80% similar across the group and generated in-silico baits from the focal genome (Figure 1B). These baits were mapped back to each member (Figure 1C) to identify single-copy orthologs within the group. Next, we designed two, overlapping 80bp baits from loci in each member of the group (Figure 1D) and then removed baits with >95% sequence similarity (Figure 1E). These steps were repeated for each pathogen group (Figure 1F). Finally, we combined baits necessary to capture 49 loci from each pathogen group into a panel that was synthesized by Daicel Arbor Biosciences.

### Museum-archived and control samples

We extracted DNA from 38 museum samples with a Qiagen DNeasy kit. Information for each specimen are provided in Table 2. Control samples were generated by spiking naïve mouse DNA with 1% parasite DNA from *Mycobacterium bovis, M. tuberculosis, Plasmodium vivax, P. falciparum*, and *Schistosoma mansoni*. An aliquot of this 1% pathogen mixture was further diluted into mouse DNA to create a 0.001% host:pathogen mixture.

**Table 2.** Specimes Examined.

| Field Accession Numbers | Museum Accession |  | Locality | Collection date | NCBI: Biosample | NCBI: SRA |
| --- | --- | --- | --- | --- | --- | --- |
|  | Number | Species |  |  |  |  |
| TK48533 | TTU75621 | *Myotis volans* | Mexico: Durango, Arroyo El Triguero | May 18, 1995 | SAMN31718202 | TDB |
| TK49668 | TTU71152 | *Didelphis virginiana* | United States: Texas, Kerr | May 14, 1996 | SAMN31718203 | TDB |
| TK49674 | TTU71137 | *Peromyscus attwateri* | United States: Texas, Kerr | May 14, 1996 | SAMN31718204 | TDB |
| TK49686 | TTU71149 | *Peromyscus laceianus* | United States: Texas, Kerr | May 14, 1996 | SAMN31718205 | TDB |
| TK49712 | TTU71215 | *Dasypus novemcinctus* | United States: Texas, Kerr | May 16, 1996 | SAMN31718206 | TDB |
| TK49732 | TTU71170 | *Lasiurus borealis* | United States: Texas, Kerr | May 17, 1996 | SAMN31718207 | TDB |
| TK49733 | TTU71171 | *Myotis velifer* | United States: Texas, Kerr | May 16, 1996 | SAMN31718208 | TDB |
| TK57832 | TTU76531 | *Peromyscus attwateri* | United States: Texas, Kerr | May 14, 1997 | SAMN31718209 | TDB |
| TK70836 | TTU81587 | *Desmodus rotundus* | Mexico: Durango, San Juan de Camarones | June 27, 1997 | SAMN31718210 | TDB |
| TK90542 | TTU131155 | *Sigmodon hirsutus* | Mexico: Chiapas, Comitán | July 9, 1999 | SAMN31718211 | TDB |
| TK93223 | TTU82694 | *Peromyscus melanophrys* | Mexico: Oaxaca, Las Minas | July 13, 2000 | SAMN31718212 | TDB |
| TK93289 | TTU82612 | *Carollia subrufa* | Mexico: Chiapas, Ocozocoautla | July 16, 2000 | SAMN31718213 | TDB |
| TK93402 | TTU82859 | *Chaetodipus eremicus* | Mexico: Coahuila | July 22, 2000 | SAMN31718214 | TDB |
| TK101275 | TTU83936 | *Glossophaga commissarisi* | Honduras: Comayagua, Playitas | July 10, 2001 | SAMN31718215 | TDB |
| TK136205 | TTU103827 | *Heteromys desmarestianus* | Honduras: Atlantida, Jardin Botanico Lancetilla | July 16, 2004 | SAMN31718216 | TDB |
| TK136222 | TTU104090 | *Peromyscus mexicanus* | Honduras: Colon, Trujillo | July 17, 2004 | SAMN31718217 | TDB |
| TK136228 | TTU104096 | *Heteromys desmarestianus* | Honduras: Colon, Trujillo | July 17, 2004 | SAMN31718218 | TDB |
| TK136240 | TTU104106 | *Glossophaga soricina* | Honduras: Colon, Trujillo | July 16, 2004 | SAMN31718219 | TDB |
| TK136756 | TTU104172 | *Eptesicus furinalis* | Honduras: Colon, Trujillo | July 17, 2004 | SAMN31718220 | TDB |
| TK136783 | TTU104199 | *Glossophaga leachii* | Honduras: Colon, Trujillo | July 17, 2004 | SAMN31718221 | TDB |
| TK148935 | TTU110359 | *Rhogeessa tumida* | Mexico: Tamaulipas, Soto la Marina | July 27, 2008 | SAMN31718222 | TDB |
| TK148943 | TTU110031 | *Myotis velifer* | Mexico: Tamaulipas, Soto la Marina | July 27, 2008 | SAMN31718223 | TDB |
| TK150290 | TTU104701 | *Balantiopteryx plicata* | Mexico: Michoacan, El Marqués | July 22, 2006 | SAMN31718224 | TDB |
| TK154677 | TTU114227 | *Gerbilliscus leucogaster* | Botswana: Ngamiland, Koanaka Hills | June 29, 2008 | SAMN31718225 | TDB |
| TK154685 | TTU114228 | *Gerbilliscus leucogaster* | Botswana: Ngamiland, Koanaka Hills | June 29, 2008 | SAMN31718226 | TDB |
| TK154687 | TTU114229 | *Gerbilliscus leucogaster* | Botswana: Ngamiland, Koanaka Hills | June 29, 2008 | SAMN31718227 | TDB |
| TK164683 | TTU115216 | *Mastomys natalensis* | Botswana: Ngamiland, Koanaka Hills | July 18, 2009 | SAMN31718228 | TDB |
| TK164686 | TTU114965 | *Mastomys natalensis* | Botswana: Ngamiland, Koanaka Hills | July 18, 2009 | SAMN31718229 | TDB |
| TK164689 | TTU114968 | *Mastomys natalensis* | Botswana: Ngamiland, Koanaka Hills | July 18, 2009 | SAMN31718230 | TDB |
| TK164690 | TTU114969 | *Mastomys natalensis* | Botswana: Ngamiland, Koanaka Hills | July 18, 2009 | SAMN31718231 | TDB |
| TK164702 | TTU114971 | *Mastomys natalensis* | Botswana: Ngamiland, Koanaka Hills | July 19, 2009 | SAMN31718232 | TDB |
| TK164714 | TTU114980 | *Mastomys natalensis* | Botswana: Ngamiland, Koanaka Hills | July 19, 2009 | SAMN31718233 | TDB |
| TK164728 | TTU114988 | *Mastomys natalensis* | Botswana: Ngamiland, Koanaka Hills | July 19, 2009 | SAMN31718234 | TDB |
| TK166246 | TTU130716 | *Peromyscus attwateri* | United States: Texas, Kerr | May 17, 2010 | SAMN31718235 | TDB |
| TK179690 | TTU137695 | *Peromyscus attwateri* | United States: Texas, Kerr | May 20, 2013 | SAMN31718236 | TDB |
| TK185677 | TTU143718 | *Peromyscus attwateri* | United States: Texas, Kerr | May 21, 2018 | SAMN31718237 | TDB |
| TK197046 | TTU143723 | *Peromyscus attwateri* | United States: Texas, Kerr | May 26, 2016 | SAMN31718238 | TDB |
| TK199855 | TTU149214 | *Peromyscus attwateri* | United States: Texas, Kerr | May 21, 2019 | SAMN31718239 | TDB |

### Library preparation

Standard DNA sequencing libraries were generated from 500 ng of DNA per sample. We combined individual libraries with similar DNA concentrations into pools of 4 samples and used the high sensitivity protocol of myBaits® v.5 (Daicel Arbor Biosciences) to enrich target loci. We used two rounds of enrichment (24 hours at 65°C). Unbound DNA was washed away and the remainder was amplified for 15 cycles, before being pooled for sequencing.

### Classifying reads

First, we generated a dataset of target loci by mapping the probes to representative and reference genomes in RefSeq v212 with BBMap v38.96 (27). For each probe, we kept the 10 best sites that mapped with ≥85% sequence identity along with 1000bp up and downstream. These sequences were combined into a database to classify individual reads with Kraken2 v2.1.1 (28) (Figure 3A). Next, we extracted pathogen reads with KrakenTools v1.2 (https://github.com/jenniferlu717/KrakenTools/; last accessed 7 Sept. 2022). These reads were assembled (Figure 3B) with the SPAdes genome assembler v3.14.1 (29) and filtered to remove low quality contigs (<100bp and <10x median coverage). Individuals with <2 contigs were removed from downstream analyses. During this time, we extracted target loci in available reference genomes (Figure 3C). Next, we identified (Figure 3D), aligned and trimmed (Figure 3E) orthologs, before concatenating them into a single alignment (Figure 3F). Finally, we generated and bootstrapped a phylogenetic tree (Figure 3G) with RaxML-NG v1.0.1 (30). These steps were repeated for each pathogen group (Figure 3H).

**Figure 3.**
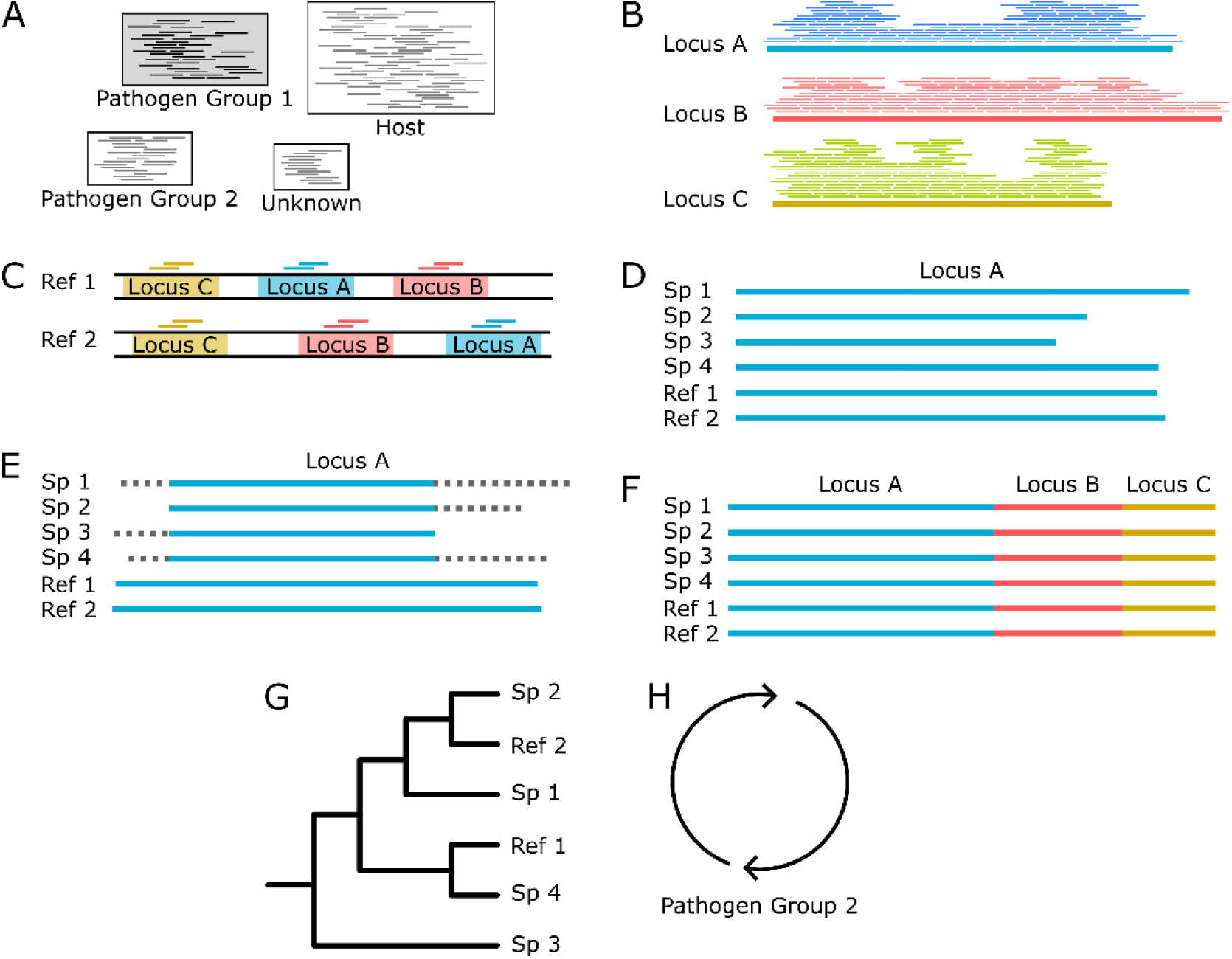
Building phylogenies from parasite reads. (A) After read classification, we extracted all the reads associated with a pathogen group. (B) These reads were then assembled into contigs with a genome assembler. (C) Simultaenously we identified and extracted the target loci from all members of the pathogen group with available reference genomes. This ensures that our final phylogeny has representatives from as many members of the pathogen group as possible. (D) Next, for each targeted locus, we combine the assembled contigs and genome extracted loci for (E) multiple sequence alignment and trimming. (F) Then, each aligned and trimmed locus is concatenated together for (G) phylogenetic analyses. (H) Finally, and if necessary, these steps are repeated for reads classified in other pathogen groups.

### Host identification

There was sufficient mtDNA sequences from most samples to verify museum identifications by comparing reads to a Kraken2 v2.1.2 (28) database of mammalian, mitochondrial genomes. We filtered the classifications by removing samples with <50 classified reads and single-read, generic classifications.

## Results

### Panel development

We used the UCE protocol developed by Faircloth *et al*. (25, 26) to develop a set of 39,893 biotinylated baits that target 32 pathogen groups responsible for 32 zoonoses. Each pathogen group is targeted at 49 loci with a few diverse taxa, *Bacillus cereus* and *Trypanosoma*, targeted at 98 loci. Table 3 contains information on pathogen groups, focal taxa, genome accession, number of baits, etc.

**Table 3.**
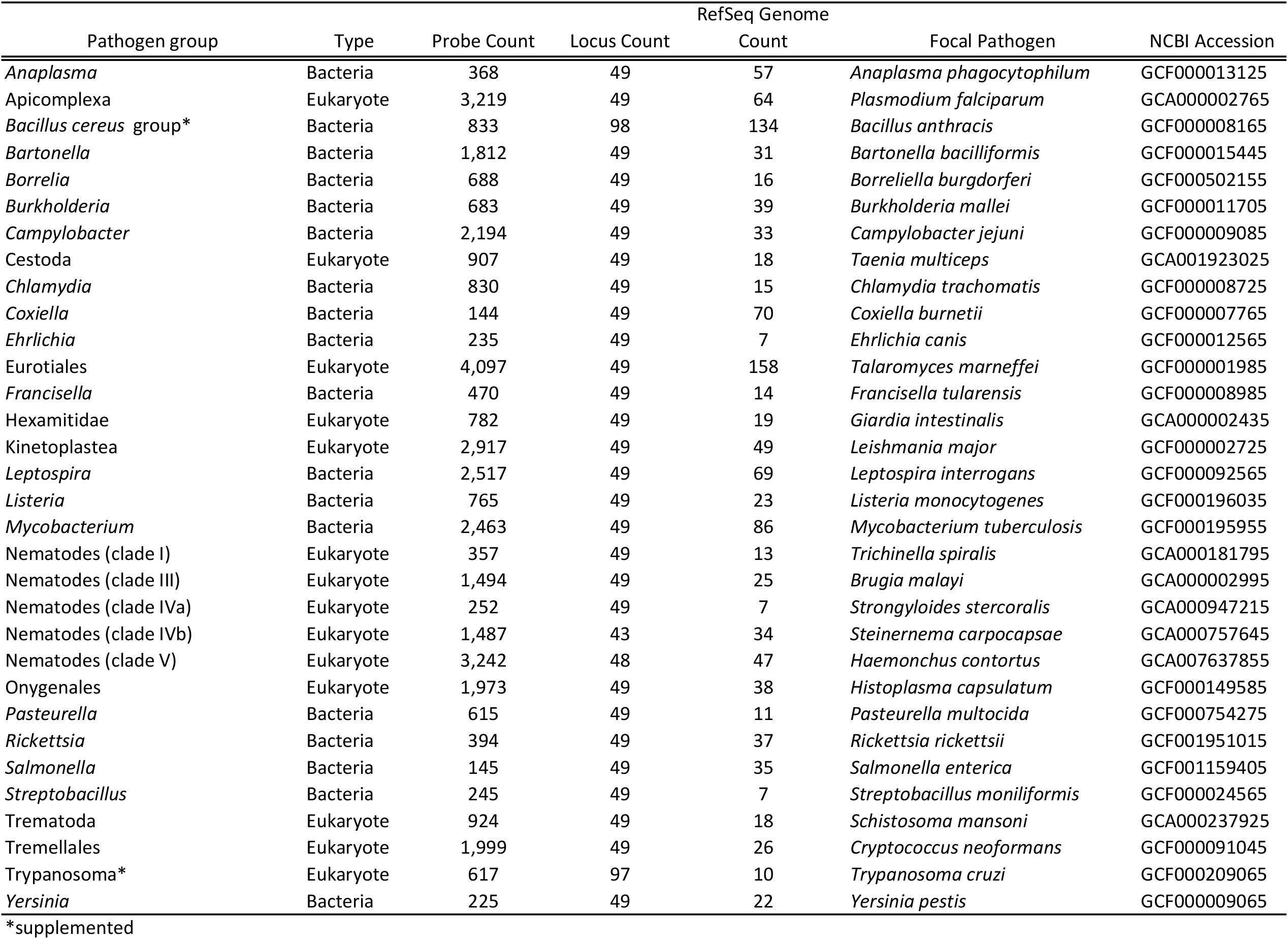
Summary of probe development.

### Control samples

We tested the efficacy of our bait set on lab-made host:pathogen mixtures of *Mus musculus, Mycobacterium tuberculosis, Plasmodium falciparum, P. vivax, and Schistosoma mansoni*. We generated four control samples containing either 1% or 0.001% pathogen DNA that was enriched or not. We classified reads against the database of target loci and found that 42.7% of all reads (*Mycobacterium* = 13.1%, *Plasmodium* = 28.1%, *Schistosoma* = 1.5%) were from control pathogens in the 1%, enriched control sample. However, only 0.03% of the corresponding 1% unenriched control was from target loci. Aside from the raw percentages, we compared the coverage of each probed region in the 1% enriched and unenriched control samples (Figure 4B-D) to understand how enrichment impacted coverage at each locus. Mean coverage per *Mycobacterium* locus increased from 0.14x to 944.5x (6746-fold enrichment), 0.53x to 1527.4x for *Plasmodium* loci (2882-fold enrichment), and 0.02x to 117.9x (5895-fold enrichment) for schistosome loci. The sequencing library from the 0.001% unenriched sample failed during the sequencing reaction so we do not have a baseline to examine enrichment in the 0.001% samples.

**Figure 4.**
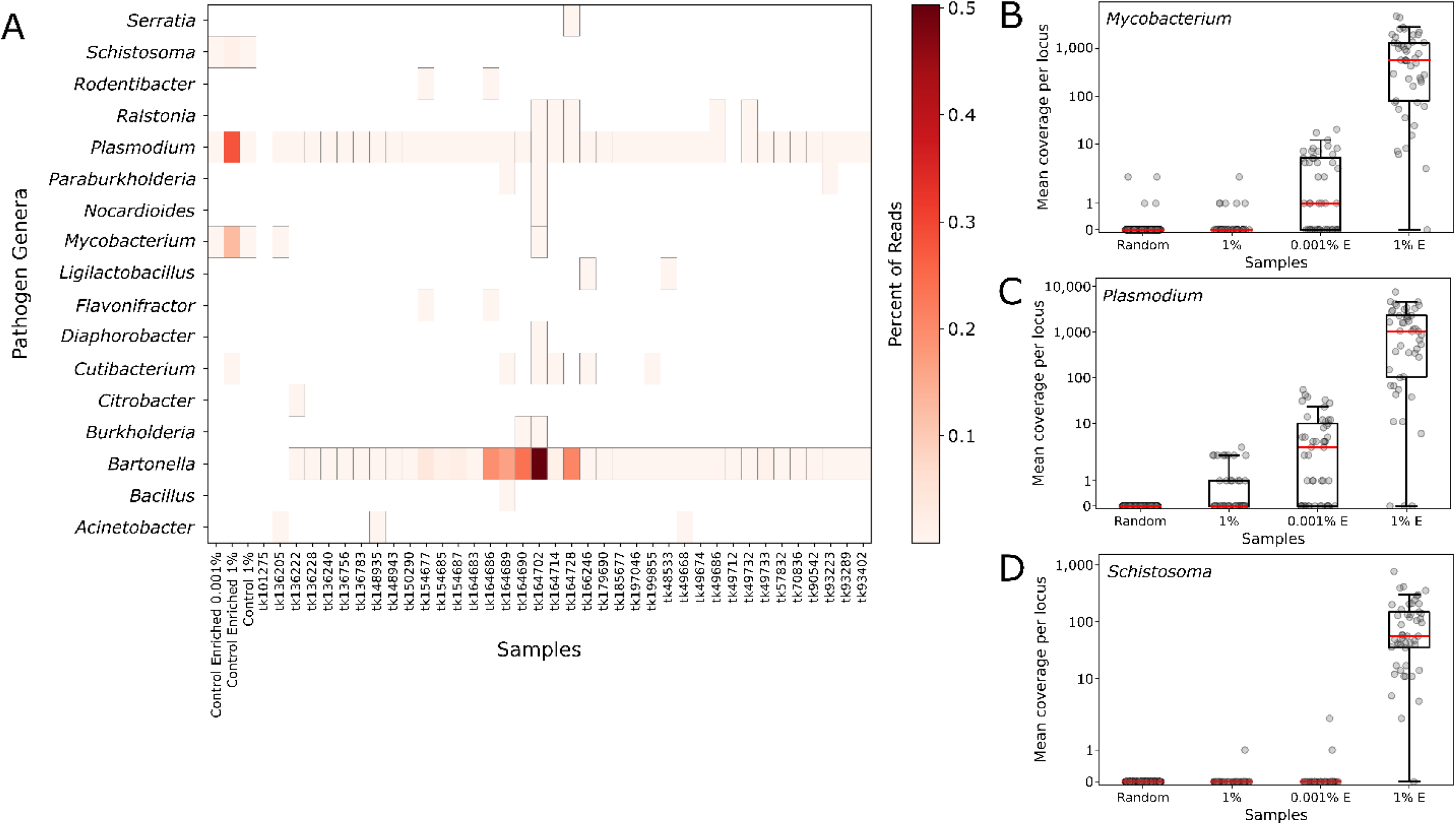
Identifying pathogen reads form controls and museum-archived tissue samples. Control reads are indicated by the percentage of pathogen DNA 1% or 0.001%. (A) Reads were compared to a database of target loci and assigned a taxonomic classification based on these results. Reads were assigned to 93 genera, of these 17 (shown) were present in at least one sample, including controls, with ≥1,000 reads. A heatmap of these results shows the relative proportion of reads assigned to each genus. Coverage at each probed locus is shown across all control samples for (B) *Mycobacterium*, (C) *Plasmodium*, and (D) *Schistosoma*. Each point in the chart is coverage calculated at a single target locus.

We extracted reads assigned to each pathogen group, assembled and aligned them with target loci extracted from reference genomes of closely related species using tools from Phyluce v1.7.1 (25, 26). We were able to assemble 0-23 target loci per pathogen group in the control samples (Table 4). Assembled loci varied in size from 109 to 1,991 bp (median 636.5bp). For each sample/group with more than two loci captured, we generated a phylogenetic tree along with other members of the taxonomic group (Figure 5). In each case, pathogen loci from the control samples were sister groups to the appropriate reference genome with strong bootstrap support. For example, the *Schistosoma* loci assembled from the 1% enriched control sample, were sister to the *Schistosoma mansoni* genome (GCA000237925) in 100% of bootstrap replicates.

**Table 4.** Parasite reads identified in and loci assembled from control samples.

| Enriched | Pathogen | Total | *Schistosoma* |  | *Plasmodium* |  | *Mycobacterium* |  |
| --- | --- | --- | --- | --- | --- | --- | --- | --- |
|  | Concentration | # Reads | # Reads | # Loci | # Reads | # Loci | # Reads | # Loci |
| TRUE | 0.001% | 509,672 | 3 | 0 | 168 | 7 | 556 | 0 |
| TRUE | 1% | 398,469 | 5,879 | 23 | 52,274 | 8 | 112,141 | 23 |
| FALSE | 1% | 375,786 | 15 | 0 | 17 | 0 | 83 | 0 |

**Figure 5.**
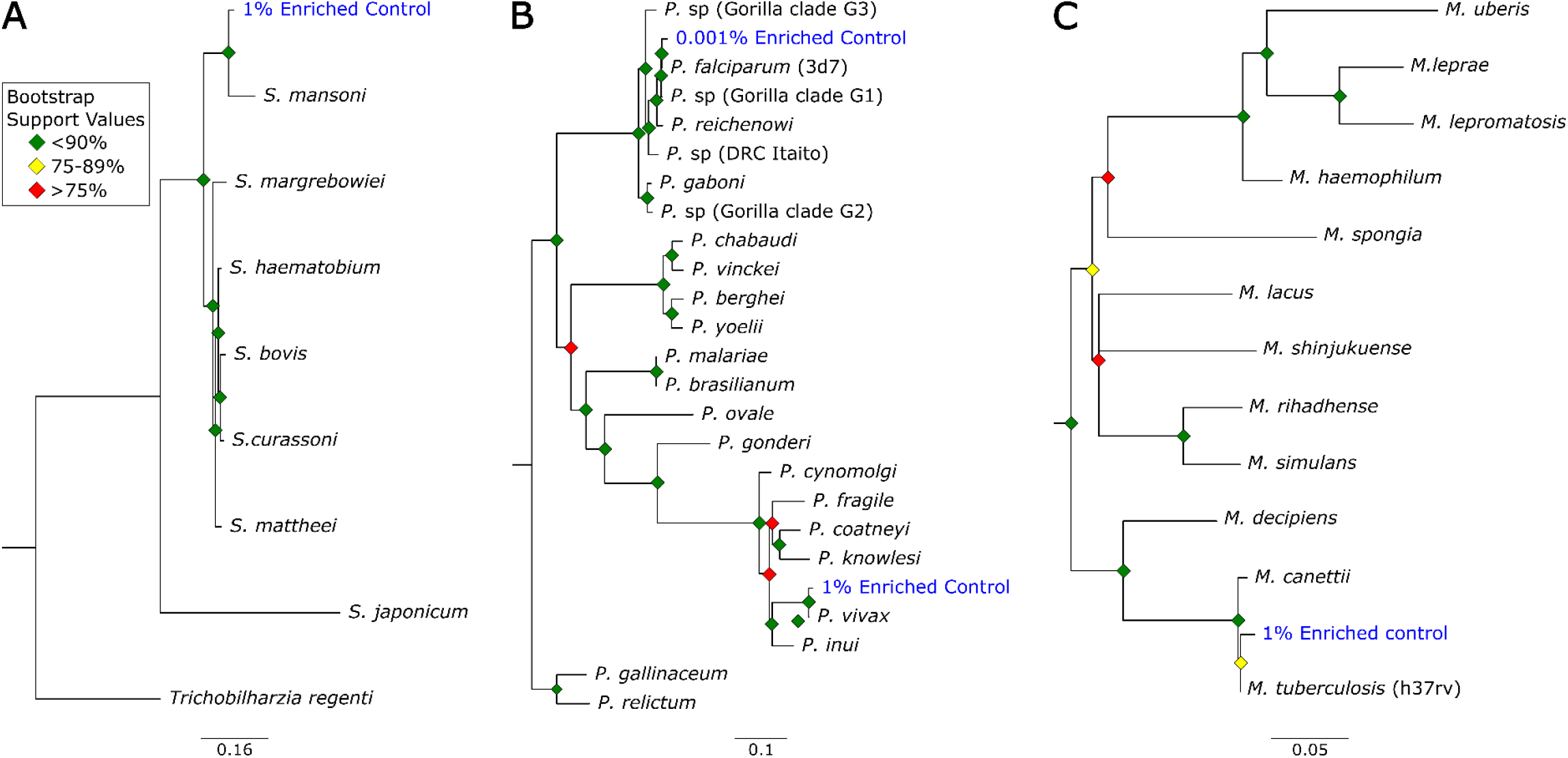
Phylogenetic analysis of pathogens using in control samples. Reads from each control pathogen (*Mycobacterium tuberculosis, Plasmodium falciparum, P. vivax, and Schistosoma mansoni*) were extracted, assembled, aligned, and trimmed for maximum likelihood, phylogenetic analyses. These phylogenies were used to identify the species or strain of pathogen used in the controls for (A) *Schistosoma*, (B) *Plasmodium*, and (C) *Mycobacterium*. Control samples are highlighted in “blue” and list the proportion of pathogen DNA. Bootstrap support values are indicated by colored “diamonds” at each available node. Branches with >50% bootstrap support were collapsed. Nodal support is indicated by color coded diamonds. Assembly accession numbers (ex. “GCA902374465”) and tree files are available as Supplemental File X.

### Museum Samples

Next, we tested our bait set on archived museum tissues. We generated 649.3 million reads across all 38 samples (mean = 17.1 million reads per sample). An initial classification showed that, on average, 4.3% of reads were assignable to loci in the database. These reads initially were designated to 93 different genera. However, 78 of those genera were at very low frequency (≤ 1,000 reads/sample (Figure 4A). Many of these low frequency hits are likely the result of bioinformatic noise (see below). *Bartonella* and *Plasmodium* were the most common genera, each present in 36 of 38 museum samples. The distribution of *Bartonella* reads was strongly bimodal such that 18 samples had ≤12 reads and 18 samples had more than 1,000 reads (median = 552). In 5 samples, the percentage of *Bartonella* reads was exceedingly high (>10%). By comparison the median number of *Plasmodium* reads never exceeded 0.04% of reads from a single museum sample (mean = 158.5 reads per sample).

We used phylogenetic analyses and rules of monophyly to identify putative pathogens to species or strain for each of the 15 genera with 1,000 or more reads (Figure 4A). We were unable to assemble more than 1 target locus for any individual in 13 genera. We were able to assemble 3-20 loci (mean = 8 loci per sample) from 16 samples containing *Bartonella* (Figure 6A), 3 loci from a sample containing *Paraburkholderia* reads (Figure 6B), and 8 loci from a sample containing *Ralstonia* reads (Figure 6C).

**Figure 6.**
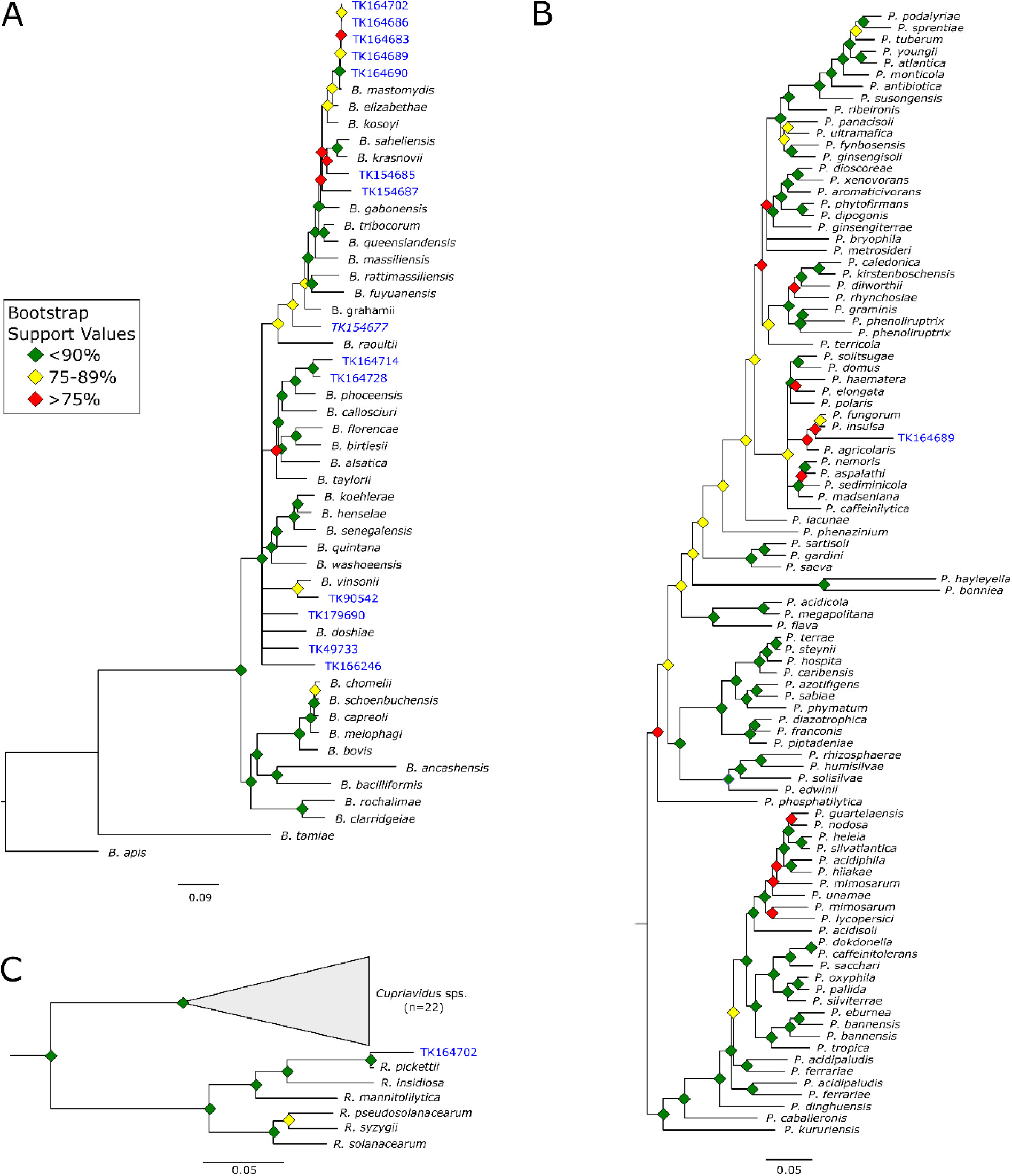
Phylogenetic analysis of pathogens using museum archived samples. These phylogenies were used to identify the species or strain of (A) *Bartonella*, (B) *Ralstonia*, and (C) *Paraburkholderia*. Museum archived samples are highlighted in “blue” and use the museum accession number (Table 1). Branches with >50% bootstrap support were collapsed. Nodal support is indicated by color coded diamonds. Assembly accession numbers (ex. “GCA902374465”) and tree files are available as Supplemental File 1.

### Host identification

We compared reads from each sample to a database of mitochondrial genomes to identify the host. In general, reads from the mitochondria comprised a small proportion (≤1%; mean=0.04%) of each sample (Figure 7). Despite the low number of mitochondrial reads, generic classifications from the mitochondrial database coincided with the museum IDs after filtering samples with ≤ 50 mitochondrial reads. For the remaining samples, the correct genus was identified by >85% of reads from that sample (mean=98%). Classifying reads below the generic level is limited by mitochondrial genome availability, but where possible, we were able to confirm museum identifications at the species level.

**Figure 7.**
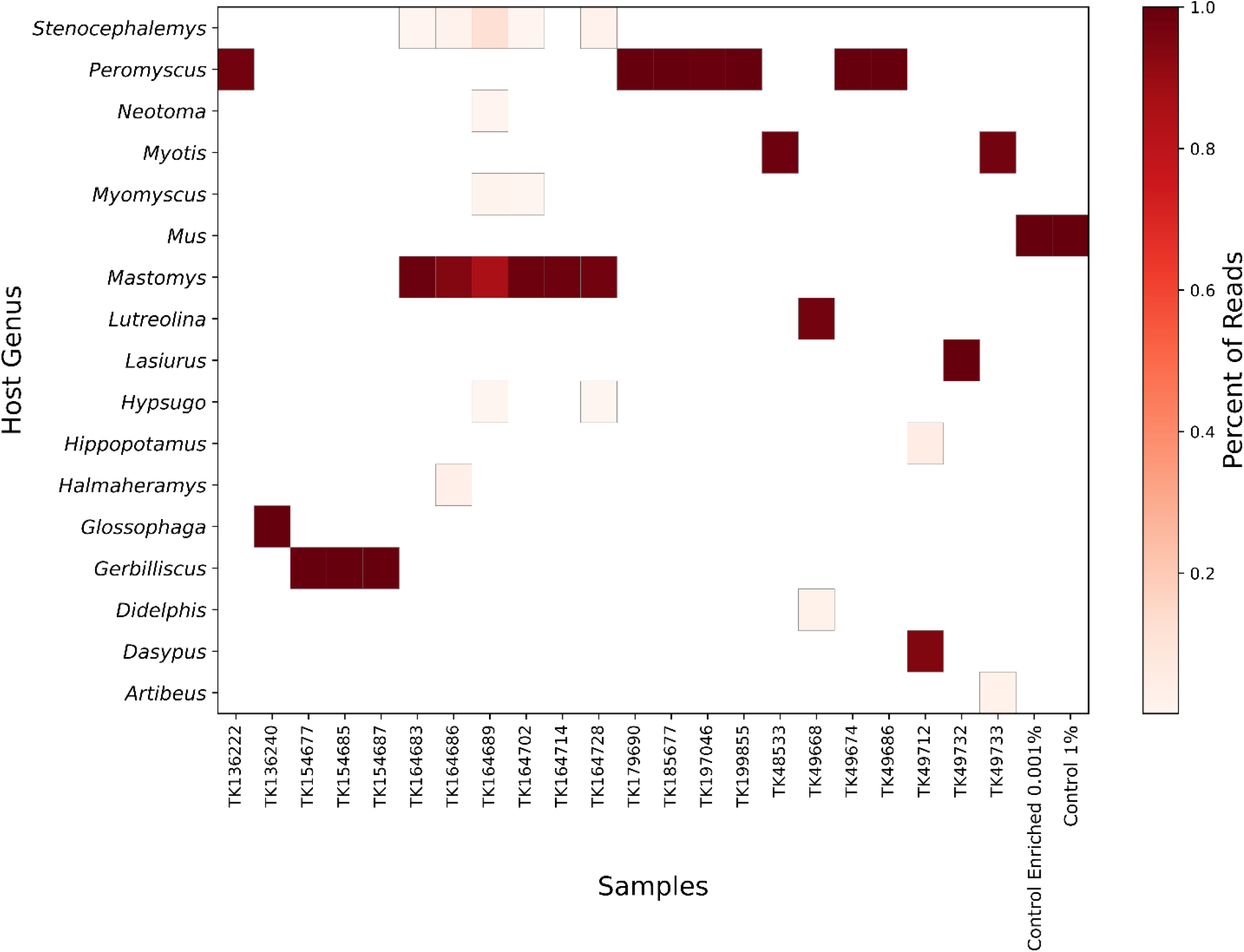
Genetic identification of host. Reads were compared to database of mammalian mitochondria and assigned a taxonomic classification based on these results. A heatmap of these results shows the relative proportion of classified reads assigned to mammalian genera. Samples with fewer than 50 mitochondrial reads and single read genera are not shown.

**Figure 8.**
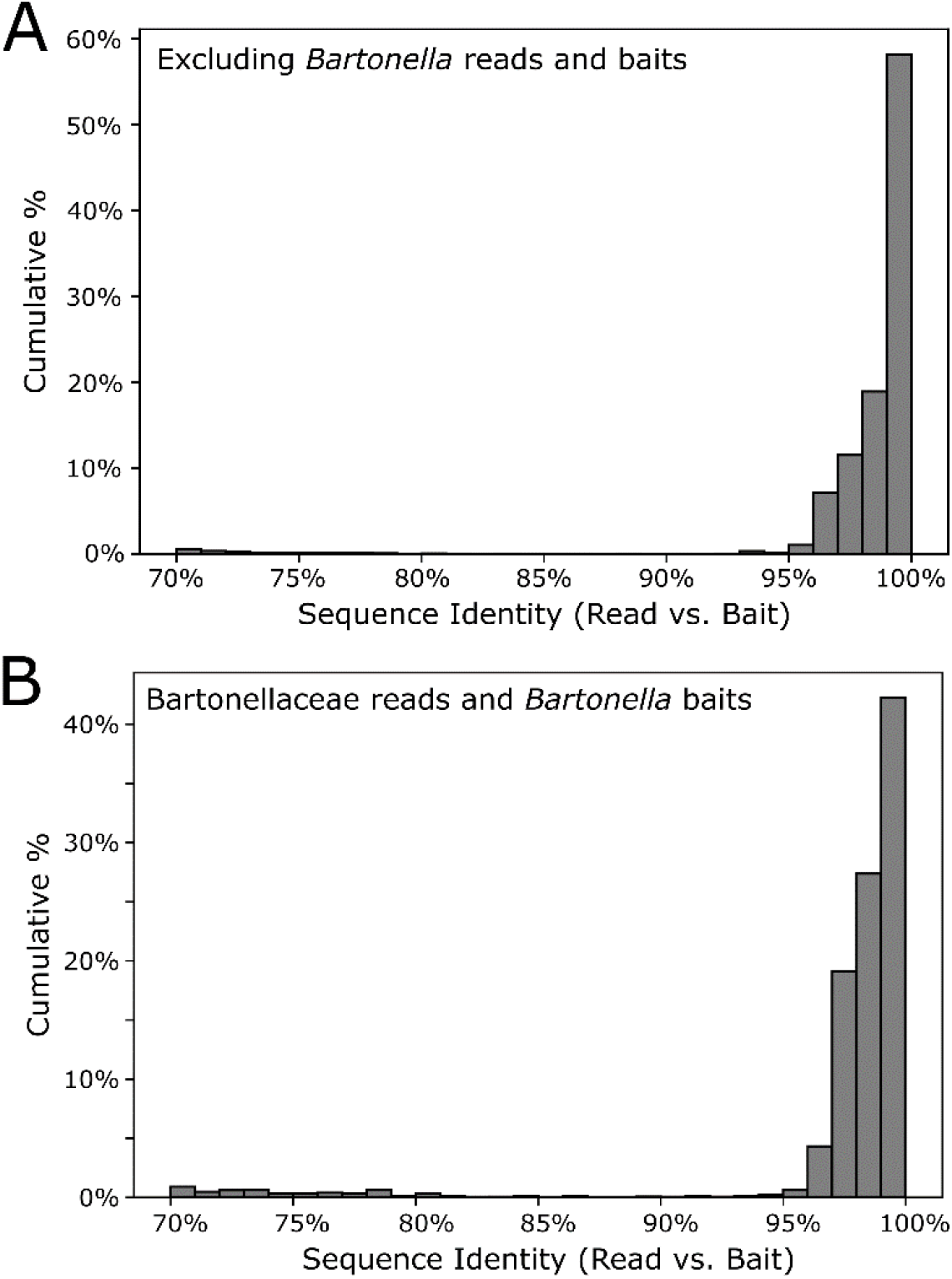
Sequence identity between enriched reads and baits. Reads from each sample were classified against a database of target loci separate host vs. pathogen derived reads. Here we show sequence identity between pathogen derived reads and the most similar bait in the bait panel for (A) all pathogens (excluding *Bartonella*). *Bartonella* was the most common pathogen in our samples and the number of reads were biased towards a few individuals. (B) *Bartonella* derived reads are shown separately.

## Discussion

We developed a set 39,893 biotinylated baits for targeted sequencing of >32 zoonotic pathogens, and their relatives, from host DNA samples. To test the efficacy of the bait panel we used four control samples that contained either 1% or 0.001% pathogen DNA and further subdivided into pools that were enriched and unenriched. Our results (Figure 4) show a large increase of pathogen DNA in the 1% enriched sample when compared to its unenriched counterpart. Specifically, enrichment increased the amount of pathogen DNA from 0.03% to 42.1%.

### Level of Detection

We were able to generate phylogenetically informative loci from *Plasmodium, Mycobacterium*, and *Schistosoma* in our 1% enriched control sample. Based on genome size we estimate 91,611, 261,030, and 3,159 copies of each genome respectively in the control sample. These results indicate the probe set is able to detect these pathogens from even a few thousand genome copies per sample (*Schistosoma*). By contrast we were only able to generate phylogenetically informative loci from *Plasmodium falciparum* in 0.001% enriched sample which would hypothetically contain ∼39 genome copies. This would imply that the bait set may be capable of identifying pathogens present in samples with a few hundred of genome copies, however there are caveats to *Plasmodium* detection that should be considered. These are discussed below.

### Enrichment at homologous loci

In each of the samples, the reads were detected from only a few loci rather than from the entire genome. For example, in the 1% enriched sample 5,879 of the 398,469 reads came from 32 loci totaling 19.6 Kb. Had the unenriched sample contained the same number of reads, randomly distributed across the genome, it would have amounted to 1 read every 62 Kb. Further, we found that enrichment increased coverage at probed loci from 0.23x to 863.3x, a 3732.3-fold averaged across all pathogens/loci (Figure 4). These results show that although large amounts of host DNA may remain in a sample, the targeted loci are significantly enriched.

### Pathogen capture from museum specimens

We tested the panel of baits on 38, museum-archived, wild, small mammal samples without any prior knowledge of infection history. Reads from these samples were initially designated to 93 different genera, but most of these were genera contained in a limited number or reads. For example, almost half of the 93 genera (n=43) were identified based on the presence of a single read across all 38 samples; most likely a bioinformatic artifact (see below). We identified 15 genera where one sample had ≥1,000 reads. For each of these 15 genera, we extracted any reads classified within the same family (ex. genus *Bartonella*, family Bartonellaceae), assembled, aligned, and trimmed them for phylogenetic analyses. In most cases the reads failed the assembly step (n=6), were filtered based on locus size or coverage (n=5), or assembled into multiple loci that were not targeted by our bait set (n=2), after which these were not pursued any further. We were, however, able to generate phylogenies for individuals positive for *Bartonella, Ralstonia*, and *Paraburkholderia*.

### Phylogenetics of pathogen taxa from sequence capture

*Bartonella* is a bacterial genus responsible for cat-scratch fever, Carrión’s disease, and trench fever (31). Transmission often occurs between humans and their pets, or from infected fleas ticks, ticks, or other arthropod vectors (32). We were able to recover target loci for 14 of 36 individuals. A phylogeny of *Bartonella* placed the museum samples in multiple clades (Figure 6A). For example, 5 individuals formed a monophyletic clade sister to *B. mastomydis. B. mastomydis* recently was described from *Mastomys erythroleucus* collected in Senegal (33). Appropriately, these samples were collected from *M. natalensis* from Botswana (Table 2). Another clade contained *B. vinsonii* and a *Sigmodon* (TK90542) collected in Mexico. Zoonotic transmission of *B. vinsonii* has been implicated in neurological disorders (34). Other museum samples likely contain novel *Bartonella* species/strains or, at the very least, they represent species/strains without genomic references.

*Paraburkholderia* is a genus of bacteria commonly associated with soil microbiomes and plant tissues. We identified *Paraburkholderia* reads in three individuals, and were able to place one of these in a phylogeny sister to a clade containing *P. fungorum* and *P. insulsa*. Bootstrap values across the phylogeny were moderate in general, and weak in this particular region (Figure 6B) so placement of this sample is tenuous. *P. fungorum* is the sole member of *Paraburkholderia* thought capable of infecting humans, but it is only a rare, opportunistic, human pathogen (35-37).

*Ralstonia* is a bacteria genus closely related to *Pseudomonas*. We identified *Ralstonia* reads in five samples, and were able to place an individual on a phylogeny. This sample is closely affiliated with *R. pickettiii* (Figure 6C). We are unaware any examples of zoonotic transmission of *R. pickettii*. Rather *R. pickettii* has been identified as a common contaminant in laboratory reagents (38) and outbreaks have been caused by contaminated medical supplies (39). We failed to identify nucleic acids in any of our negative controls during library preparation. Further, if there were systemic contamination we would expect to find *Ralstonia* in all of our samples rather than the 5 of 36 observed here. That said, because we cannot rule out reagent contamination, the presence of *Ralstonia* in the museum samples should be interpreted with caution.

### Efficient capture of diverse pathogens species from target groups

We were able to capture, sequence, and assemble loci from taxa that were not represented in the databases used to design the bait panel. This was possible for two reasons. First, the bait panel is highly redundant. The baits themselves are sticky and able to capture nucleic acid fragments ≤10-12% diverged (40). We designed the panel with ≤5% sequence divergence between any pair of baits at a particular locus. Second, sampled loci within each pathogen group spanned a range of divergences. Conserved loci were more likely to catch more divergent species that may not have been present in our initial dataset. For example, we catch multiple species of Bartonella that were not present in our probe set, for which related genomes were available. For *Ralstonia* and *Paraburkholderia*, however, we identified these samples from reads targeted by probes for Burkholderia a pathogenic taxon in the same family (Burkholderacea). The ability to identify taxa at these distances is because of the more conserved loci targeted by the bait panel.

### Widespread *Plasmodium* Detection

During the initial read classification stage, we identified low levels of *Plasmodium* in all but two museum samples, which was unexpected. Museum samples contained ≤3,221 *Plasmodium* reads per sample (mean = 428.3 reads) but we were unable to assemble them into loci for phylogenetic analyses. This effectively removed these samples from downstream analyses. The *P. falciparum* genome is extremely AT-rich (82%; 41) which may result in a bioinformatic false positives. We suspect that AT-rich, low-complexity regions of the host genome are misclassified as parasite reads. To test this hypothesis, we used fqtrim 0.9.7 (https://ccb.jhu.edu/software/fqtrim) to identify and remove low complexity sequences within these reads. This filter, by itself reduced the number of *Plasmodium* reads in the museum samples by 75.5% (max = 298 reads, mean = 57.2 reads). By comparison, only 8.2% and 0.2% of the reads from 0.001% and 1% enriched control samples were removed.

### Challenges and future prospects

Several technical issues still need to be addressed. First, enrichment increases the targeted loci coverage by three orders of magnitude, however, the amount of host DNA remaining in each sample is still high. Ideally, host DNA would be rare or absent. Second, the bait panel itself requires relatively large up-front costs. Third, although the bait panel is developed to target a wide range of taxa it is not possible to know which species are missed. The best way to circumvent this issue is to use controls spiked with various pathogens of interest similar to how mock communities are used in other metagenomic studies (42). These mock controls are commercially available for bacterial communities (ex. ZymoBIOMICS Microbial Community Standards, Zymo Research), but we have been unable to find similar products that contain eukaryotic pathogens. Solutions to these problems will make targeted sequencing with bait panels a viable tool for pathogen surveillance. Fourth, the sensitivity of the probes will depend on the sequence divergence between the probes and pathogen DNA. The more diverged the two are, the less efficient the capture will be. This means that pathogen groups with biased or limited genomic data will be less likely to capture off-target species once divergence increases by more than 5-10%. Finally, the current probe panel is capable of capturing and identifying pathogens if there are ≥3,000 genome copies in the sample. Sensitivity needs to be improved in future iterations of the panel. One method could be to target pathogen specific, repetitive sequences (43). Since these sequences are already present in the genome hundreds to thousands of times, it should be possible to significantly increase the sensitivity of the probe panel.

Although further effort is required to resolve these issues, we believe that enrichment of pathogen DNA from museum tissue samples is a viable tool worth further development. At the very least, in its current form, it represents a coarse tool that can be used to “scan” for the presence of various pathogens from archived tissues. Tools like target enrichment will be necessary for maximizing the pathogen data that is available from the hundreds of thousands of museum-archived tissues and will play a critical role in understanding our susceptibility to future zoonotic outbreaks.

## Supporting information

Supplemental Tree File

## Acknowledgements

We thank the following people: Sandy Smith, John Heaner, Larry Schlesinger, Ian Cheeseman, and Frederic Chevalier from Texas Biomedical Research Institute for providing computational and laboratory support. Kathy McDonald, Heath Garner, and Caleb Phillips at the Natural Science Research Laboratory at Texas Tech University provided small mammal tissues. This research was funded by the Texas Biomedical Research Forum (19-04773).

## Supplemental Materials and Methods

### Host-Pathogen control sample

We isolated DNA using the Qiagen blood and tissue kit following manufacturer’s protocol and quantified DNA using Qubit. We prepared a cocktail of pathogen DNA mixtures comprising 200 ng DNA of each pathogen (*Mycobacterium bovis, M. tuberculosis, Plasmodium vivax, P. falciparum, Schistosoma mansoni*, and *S. bovis*). A mammalian-pathogen DNA mixture was prepared by mixing pathogen DNA in DNA from uninfected liver tissues of laboratory mouse (*Mus musculus*) to make 1% and 0.001% pathogen mixtures. The negative control was prepared from the same uninfected liver tissues of laboratory mouse (*Mus musculus*) without spiking with pathogens.

### Museum-archived samples and controls

We extracted DNA from 42 museum samples comprising of mammalian liver tissues (in lysed buffer or frozen in liquid nitrogen) collected between 1995 and 2018 in Africa, Southern America and the USA. Control samples were as previously described (1% and 0.001% pathogens DNA in mammalian DNA). Information for each specimen are provided in Table 2.

### Computing environment and reproducibility

All analyses were performed on a single compute node with 48 processors and limited to 100Gb of RAM. Bioinformatic steps were documented in a series of BASH shell scripts or Jupyter v4.9.2 notebooks. These files along with conda v4.11.0 environments are available at github.com/nealplatt/pathogen_probes and archived at (DOI: 10.5281/zenodo.7319915).

### Panel development

We developed a set of biotinylated probes for UCE-based, targeted sequencing of 32 pathogen groups (Table 1). For example, given the large evolutionary distances covered by various pathogens, we generated sets of probes that target more discrete taxonomic groups (ex. Nemotoda, *Yersinia*, etc). For bacterial pathogens, probes were designed to capture all species within the genus or species group. For eukaryotic pathogens, probes were designed to be effective at taxonomic ranks that ranged from species group to class. The taxonomic rank varied in eukaryotic pathogens based on the following criteria: 1) the number of available genomes, 2) sequence diversity - because this impacted the number of probes needed. Table 1 provides information on the pathogen group, targeted zoonotic agent and zoonoses.

For each group we used the Phyluce package v1.7.0; (1, 2) we generated probes to target ∼49 loci using the methods described below. First, we identified orthologous loci between a focal pathogen and the remaining species in the pathogen group. Focal taxa were chosen based on their assembly contiguity or prominence as a zoonotic agent. To do this we downloaded a genome for each species in the pathogen group. Accession numbers for these assemblies are provided in Table 2. Next, we simulated 25x read coverage for each genome using the ART v2016.06.05; (3) read simulator with the following options “art_illumina --paired --len 100 --fcov 25 --mflen 200 --sdev 150 -ir 0.0 -ir2 0.0 -dr 0.0 -dr2 0.0 -qs 100 -qs2 100 -na”. Simulated reads from all query taxa were mapped back to a focal taxon with bbmap v38.93; (4) allowing up to 10% sequence divergence (“minid=0.9”). Unmapped, or multi-mapping reads were removed using Bedtools v2.9.2 (5) and phyluce_probe_strip_masked_loci_from_set (filter_mask 25%). The remaining reads were merged to generate a BED file containing orthologous regions between the query and focal taxa.

Then, we identified orthologous loci among all taxa within the pathogen group using phyluce_probe_query_multi_merge_table. Next, we filtered each set of loci to retain only those shared among 33% of taxa in the pathogen group using phyluce_probe_query_multi_merge_table. We extracted 160 bp from each locus and generated an initial set of in-silico probes directly from the focal genome using phyluce_probe_get_genome_sequences_from_bed and phyluce_probe_get_tiled_probes. Additional options for probe design included generating two probes per locus (“-two_probes”) that overlapped in the middle (-overlap-middle). Focal probes with repetitive regions or skewed GC content (<30% or >70%) content were removed. Next, the probes from the focal taxa were mapped back to each genome in the pathogen group with phyluce_probe_run_multiple_lastzs_sqlite. We used the “--identity” option to limit searches with a maximum divergence of 30%. Using these results, we extracted 120 bp loci from the probed regions in each representative genome extracted using phyluce_probe_slice_sequence_from_genomes. Theoretically, this dataset should contain orthologous 120 bp sequences from most taxa in each pathogen group. We verified this with phyluce_probe_get_multi_fasta_table which provides a table showing the number of taxa identified at each locus. We used this information to identify the 100 loci capable of capturing most taxa from the pathogen group. Next, we generated two 80bp probes from each of the 100, 120 bp loci. We used phyluce_probe_easy_lastz to compare the probes to themselves and remove any that were possible duplicates. Then we reduced the probe set even further by clustering probes based on sequence identity with cd-hit-est v4.8.1;(6). We identified sequence clusters with >95% similarity and retained only one probe per group. Finally, we re-calculated the number of probes needed to capture each locus.

The proceeding steps were repeated for each pathogen group in Table 1. To generate a final panel, we selected 49 loci per pathogen group in a way that minimized the number of probes needed. In some cases, we needed to generate two sets of probes to adequately represent target pathogens. For example, Kinetoplastea contains two pathogens of interest, *Trypanosoma* and *Leishmania*. The baits designed for *Leishmania* were able to target all 49 loci in the most of the Kinetoplastea but only 23 loci in *Trypanosoma*. We then generated a second set of 617 *Trypanosoma-*specific baits to augment the kinetoplastid baits and ensure that *Trypanosoma* parasites were represented adequately in the final panel. Likewise, we doubled the number of baits used to capture loci from the *Bacillus cereus* group to effective capture *B. cereus* and *B. anthracis*. The final set of probes were quality-checked and synthesized by Arbor Biosciences.

### Library preparation

Standard DNA sequencing libraries were generated from 500 ng of DNA per sample. We used the KAPA Hyperplus kit protocol with the following modifications: i) enzymatic fragmentation at 37°C for 10 minutes, ii) adapter ligation at 20°C for an hour, and iii) 4 cycles of library PCR amplification. To minimize adapter switching we used unique dual indexed (UDI) adapters (IDT xGen Stubby Adapter-UDI Primers). Each library was eluted in 20 μL of sterile water and the base pairs sizes and concentration estimated by Agilent 4200 Tapestation® (Main Document Figure 2).

Individual samples with similar DNA concentrations were combined together into pools of 4-16 samples and the total volume was reduced to 7 μL with a speedvac vacuum concentrator. Next, we used the high sensitivity protocol of myBaits® v.5 (Daicel Arbor Biosciences) to enrich target pathogen loci from the host/pathogen control and museum archived samples. We used two rounds of enrichment for each pool of samples.Probe concentration was 100ng/μL. Each round was 24 hours at 65°C. After washing of unbound DNA, each library was amplified with a 15 cycle PCR amplification step and quantified using qPCR. Finally, the pools of 4-16 were combined into an equimolar pool for sequencing. All sequencing reactions were on single lanes of Illumina Hi-Seq 2500.

### Bioinformatic analyses

All analyses were performed on a single compute node with 48 processors and limited to 100Gb of RAM. Bioinformatic steps were documented in a series of BASH shell scripts or Jupyter notebooks. These files along with conda environments are available at github.com/nealplatt/pathogen_probes and archived at (DOI: 10.5281/zenodo.7319915). The basic structure of the bioinformatic analyses are shown in Figure 3. In general, we used the Kraken2 v2.1.2 (7) to assign a taxonomic id to each read, the Phyluce v1.7.1 (1, 2) pipeline to identify, assemble, and align loci, and RaxML-NG v1.0.1 to generate phylogenies from each pathogen group of interest.

First, we used Trimmomatic v0.39 (8) to trim and quality filter low quality bases and Illumina adapters. Then, we used Kraken2 v2.1.1 (7) to compare each read from our samples to a reduced dataset of target loci using a “–conf” cutoff of 0.2. We decided to compare our reads to a reduced dataset of target loci to minimize the computational expense of these comparison. To generate the reduced database of bait-targeted loci, we downloaded one representative or reference genome from all species in RefSeq v212 (9) with genome_updater.sh v0.5.1 (https://github.com/pirovc/genome_updater; accessed 7 Sept. 2022). Then we used BBMap v38.96 (4) to map all the baits to each genome and a kept the 10 best sites that mapped with ≥85% sequence identity. Next, we extracted these hits along with 1000bp up and downstream. These sequences were combined into a single fasta file that should contain the major mapping locations for our baits.

Once reads were classified we identified genera that were known pathogens or were present in at least one sample with more than 1,000 reads. Next, we extracted reads from the relevant family with KrakenTools v1.2 (https://github.com/jenniferlu717/KrakenTools/; last accessed 7 Sept. 2022). These reads were then assembled (Figure 3B) with the SPAdes genome assembler v3.14.1 (10) using the phyluce_assembly_assemblo_spades wrapper script. We filtered out low quality contigs based on size (<100bp) and median coverage (<10x) as calculated by the SPAdes genome assembler. Next, we filtered individuals even further by removing individuals with fewer <2 contigs.

While we were assembling and filtering contigs from each isolated target loci from species with available genome assemblies, we used genome_updater.sh v0.5.1 (https://github.com/pirovc/genome_updater; accessed 7 Sept. 2022) to download one (“-A 1”) reference or representative (‘-c “reference genome”,”representative genome”) genome from either refseq or genbank (“-d refseq,genbank”) for the pathogen group. We also included at least one individual from an outlier genus to root downstream analyses. These genomes were converted to twobit format with faToTwoBit. Next, we used phyluce_probe_run_multiple_lastzs_sqlite to compare probes from the pathogen group to the genome assemblies with an identity cut off of 85% (--identity 0.85). These loci plus 1Kb of flanking sequence (“--flank 1000”) were extracted from the genome using phyluce_probe_slice_sequence_from_genomes. After extraction, the sliced loci were identified and counted using phyluce_assembly_match_contigs_to_probes (“--min-identity 90”) and phyluce_assembly_get_match_counts. Next, we combined the loci generated from our samples with those from representative and reference genomes and aligned them with phyluce_align_seqcap_align. The resulting alignments were trimmed with gblocks v0.91b (11) and phyluce_align_get_gblocks_trimmed_alignments_from_untrimmed. We then counted the number of taxa per locus alignment (phyluce_align_get_taxon_locus_counts_in_alignments) and removed taxa with fewer than 2 loci (phyluce_align_extract_taxa_from_alignments). Then we removed any loci that contain fewer than half of the expected number of taxa with phyluce_align_get_only_loci_with_min_taxa and concatenated the remaining loci into a single phylip alignment (phyluce_align_concatenate_alignments).

We used RaxML-NG v1.0.1 (12) to generate a maximum likelihood phylogenetic tree from the concatenated alignment. We ran 100 parsimony tree searches and then another 1000 replicates using the GTR+G substitution model. Branches with less than 50% support were collapsed with the Newick Utilities v1.6 (13), Newick editor (nw_ed <input_tree_file> ‘i & b<=50’). These steps were then repeated with other pathogen groups identified in the samples.

### Host identification

We verified museum identifications by comparing reads to a second Kraken2 v2.1.2 (7) database containing mammalian mitochondrial genomes. To do this, we downloaded all available mammalian mitochondrial genomes (n=1,651) from https://www.ncbi.nlm.nih.gov/genome/organelle/ (last accessed 3 November 2022). We then created a custom database and compared each of our samples to using Kraken2 and no confidence cutoffs. The Kraken2 classifications were filtered by removing any samples with fewer than 50 classified reads and any single-read, generic classifications.

## Supplemental Figures

**Supplemental Figure 1.**
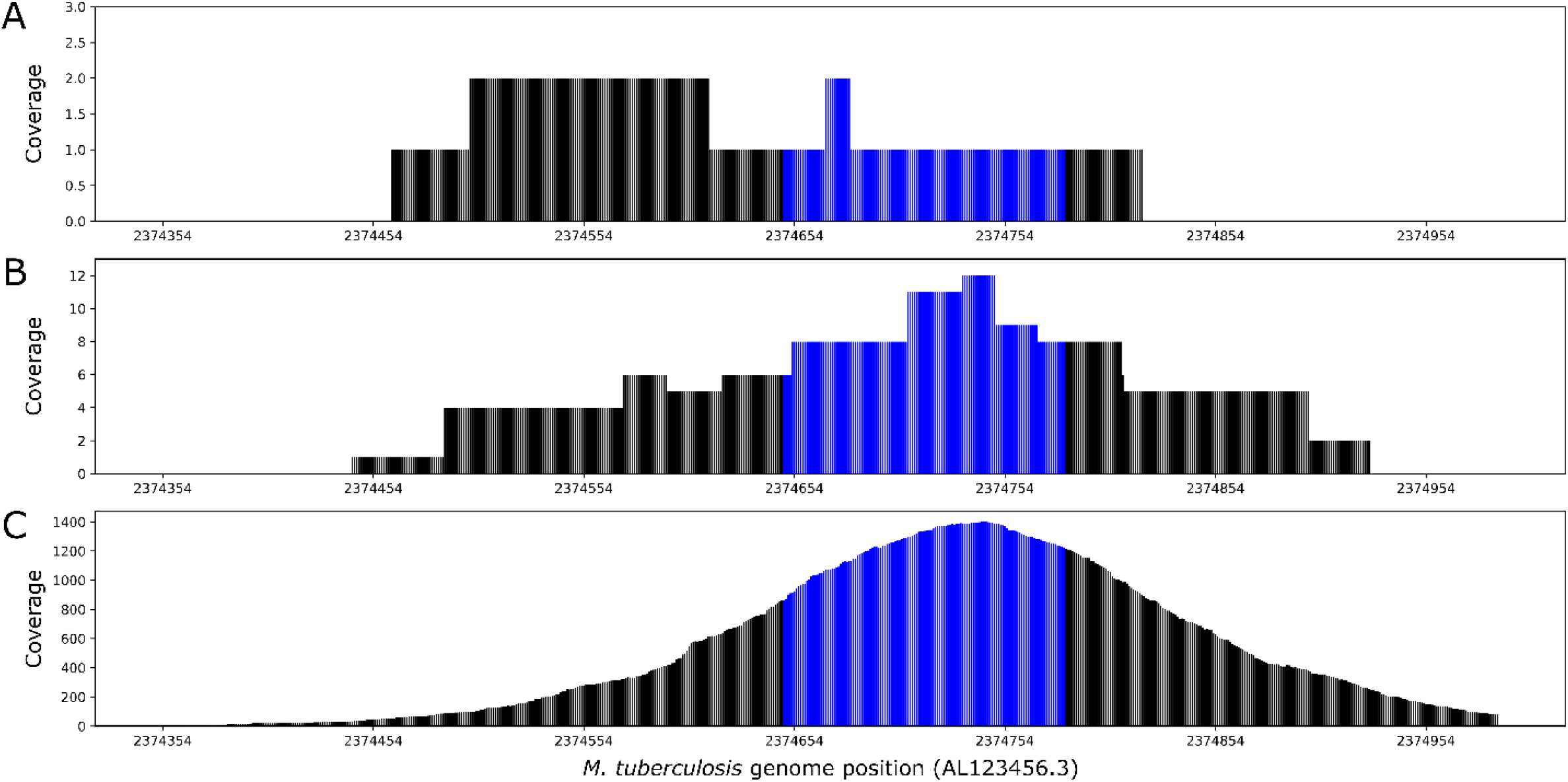
Read depth at a targeted region in *M. tuberculosis* in the (A) 1%, unenriched, (B) 0.001% enriched, and (C) 1% enriched control, samples. This particular probe was designed for (AL123456.3:2,374,648-2,374,781; shown in “blue”). Median coverage at this locus increased from 1x in the (A) 1%, unenriched to 8x in the (B) 0.001% enriched and 1278x in the (C) 1% enriched control, sample.

## Notes

### Competing Interest Statement

The authors have declared no competing interest.

